# Peptriever: A Bi-Encoder approach for large-scale protein-peptide binding search

**DOI:** 10.1101/2023.07.13.548811

**Authors:** Roni Gurvich, Gal Markel, Ziaurrehman Tanoli, Tomer Meirson

**Affiliations:** Davidoff Cancer Center, Rabin Medical Center-Beilinson Hospital, Petah Tikva, 49100, Israel; Institute for Molecular Medicine Finland (FIMM), HiLIFE, University of Helsinki, Finland; Faculty of Medicine, Tel Aviv University, Tel-Aviv, 6997801, Israel

## Abstract

**Motivation:** In peptide therapeutics, the successful interaction between a designed peptide and a specific receptor is crucial, while minimizing interactions with other receptors is equally essential. Current computational methods excel at estimating the probability of the former but estimating the latter requires excessive computational resources, making it challenging.

**Results:** In this study, we propose transformers-based protein embeddings that can quickly identify and rank millions of interacting proteins. Furthermore, the proposed approach outperforms existing sequence- and structure-based methods, with a mean AUC-ROC and AUC-PR of 0.73.

**Availability:** Training data, scripts, and fine-tuned parameters are available at https://github.com/RoniGurvich/Peptriever. A live demonstration of the application can be found at https://peptriever.app/.

## 1 Introduction

Peptides are short chains of amino acids linked by peptide bonds. A polypeptide is a long, continuous, unbranched peptide chain. Polypeptides that have a molecular mass greater than 10,000 Daltons are called proteins. Protein-peptide interactions play a vital role in numerous fundamental cellular processes, making their identification essential for peptide therapeutics(Lei et al. 2021), biosensing

(Caporale et al. 2021), and vaccine design (Di Natale et al. 2020). Due to peptide flexibility and the transient nature of protein-peptide interactions, peptides are difficult to study experimentally (Johansson-Åkhe, Mirabello and Wallner 2019). Therefore, computational methods have emerged as alternate tools that can unlock the potential of peptides as novel medicinal and biological agents (Audie and Swanson 2013). Several computational methods have been developed to predict protein-protein and protein-peptide interactions; some notable examples include InterPep2 (Johansson-Åkhe, Mirabello and Wallner 2020), PIPER-FlexPepDock(PFPD) (Alam et al. 2017), CABS-dock(Kurcinski et al. 2015) and CAMP(Lei et al. 2021). AlphaFold-Multimer, an extension of AlphaFold2, performs better than state-of-the-art methods for protein-peptide complex modeling and interaction prediction (Evans et al. 2021, Ko and Lee 2021, Johansson-Åkhe and Wallner 2022). The existing methods primarily estimate the binding probability between protein and peptide pairs. However, given a specific peptide sequence without a corresponding protein, it is challenging to identify proteins it binds with as there may be millions of candidates.

By carefully examining the pros and cons of existing methods, we developed Peptriever, a user-friendly web application that can perform peptide binding searches on millions of proteins within seconds. Peptriever utilizes an approximate nearest neighbor index populated with high-dimensional vector representations of the proteins. These vector representations, also called embeddings, are extracted using two transformers jointly trained for this purpose. Details about Peptriever’s architecture, training data, and results are discussed in the following sections.

## 2 Methods

### 2.1. Architecture and training

Peptriever employed a BERT-like architecture (Devlin et al. 2018) coupled with a byte-pair encoding tokenizer (Sennrich, Haddow and Birch 2015) to train a binding embedding model. The training process comprised two stages: unsupervised pre-training and supervised fine-tuning.

In the first stage, two separate models were trained - one for proteins and another for peptides. Following the footsteps of Protein BERT(Brandes et al. 2022), the models were trained using a Masked Language Modeling (MLM) task(Sennrich, Haddow and Birch 2015). During training, the model receives a tokenized input sequence for each sample batch with a randomly chosen subset of masked tokens. Subsequently, the model learns to replace these masked tokens with their original counterparts, leveraging information from the non-masked tokens on both sides of the sequence. Prior to the pre-training, the tokenizer was trained on the same dataset with its vocabulary size set to 1024.

It has been shown that two transformer models jointly trained with a contrastive loss function can learn to map different inputs to the same embedding space even when the inputs come from different modalities, such as text and image (Radford et al. 2021). Similar architecture called Bi-Encoder (Jung, Choi and Rhee 2022) was presented for neural ranking to improve text search.

Therefore, the two pre-trained models were jointly fine-tuned in the second stage in a similar way; the fine-tuning process aimed to minimize the distance in the embedding space between matched protein-peptide sets while maximizing the distance between a protein and all non-matching peptides. Additionally, the MLM loss was added to enhance the generalization of the embeddings.

### 2.2. Training and validation

To conduct unsupervised pre-training, 202,738 PDB files were obtained from the RCSB Protein Data Bank (Berman et al. 2000), and protein sequences were extracted into a dataset. The dataset included 40,105 genes with 159,907 unique sequences from 6,768 species.

To enable supervised fine-tuning, we integrated several databases of protein-peptide interactions across multiple species. Our data was sourced from PepBDB (Wen et al. 2019), Propedia (Martins et al. 2021), and YAPP-Cd(Park 2021), all of which are based on experimentally verified interactions with 3D structures. After removing duplicates and discarding structure information, the resulting training dataset consisted of 16,370 unique pairs of protein-peptide sequences.

To prevent data leakage caused by near-identical sequences in different training partitions, we used the gene association of the protein and the peptide to assign these to the train, validation, and test partitions. For instance, if a protein is associated with gene A and a peptide is associated with gene B, any other combinations of protein-peptide pairs in the training set sharing the same gene A and gene B associations will be grouped within the same training partition.

For evaluation, we used the test set utilized by Johansson-Akhe et al., (Johansson-Åkhe and Wallner 2022). To avoid data leaks, overlapping protein-peptide sequence sets from the test set were excluded during fine-tuning.

#### Evaluation

The quality of the embeddings was benchmarked against CAMP, InterPep2, and various versions of AlphaFold-Multimer, as previously reported in (Johansson-Åkhe and Wallner 2022). The test set defines protein-peptide interaction prediction as a binary classification task comprising 112 positive and 560 negative interactions. The performance of the model was evaluated by computing the area under the Precision-Recall curve (AUP-PR) and the area under the Receiver Operating Characteristic curve (AUC-ROC). The negated and normalized distance between the embeddings of each pair was used as the predicted probability, as shown in the following equation.

## 3 Results

The performance comparison between the proposed Peptriever and other methods is shown in **Figure 1**. Peptriever embedding achieved a higher AUC-ROC of 0.83, compared with the best-performing version of AlphaFold-Multimer (v9) with 0.81, CAMP with 0.73, and InterPep2 with 0.64. Furthermore, the model’s AUC-PR of 0.63 outperformed all versions of AlphaFold-Multimer, except v8 and v9, which achieved comparable scores of 0.63 and 0.64, respectively. It also surpassed the scores of CAMP with 0.41 and InterPep2 with 0.46. Collectively, the Peptriever embeddings exhibited a higher mean AUC-ROC and AUC-PR of 0.73 compared to all other models, including AlphaFold-Multimer v9, with a mean of 0.725. Performance improvement is possibly due to BERT-based architecture combined with a contrastive loss function.

**Figure 1.**
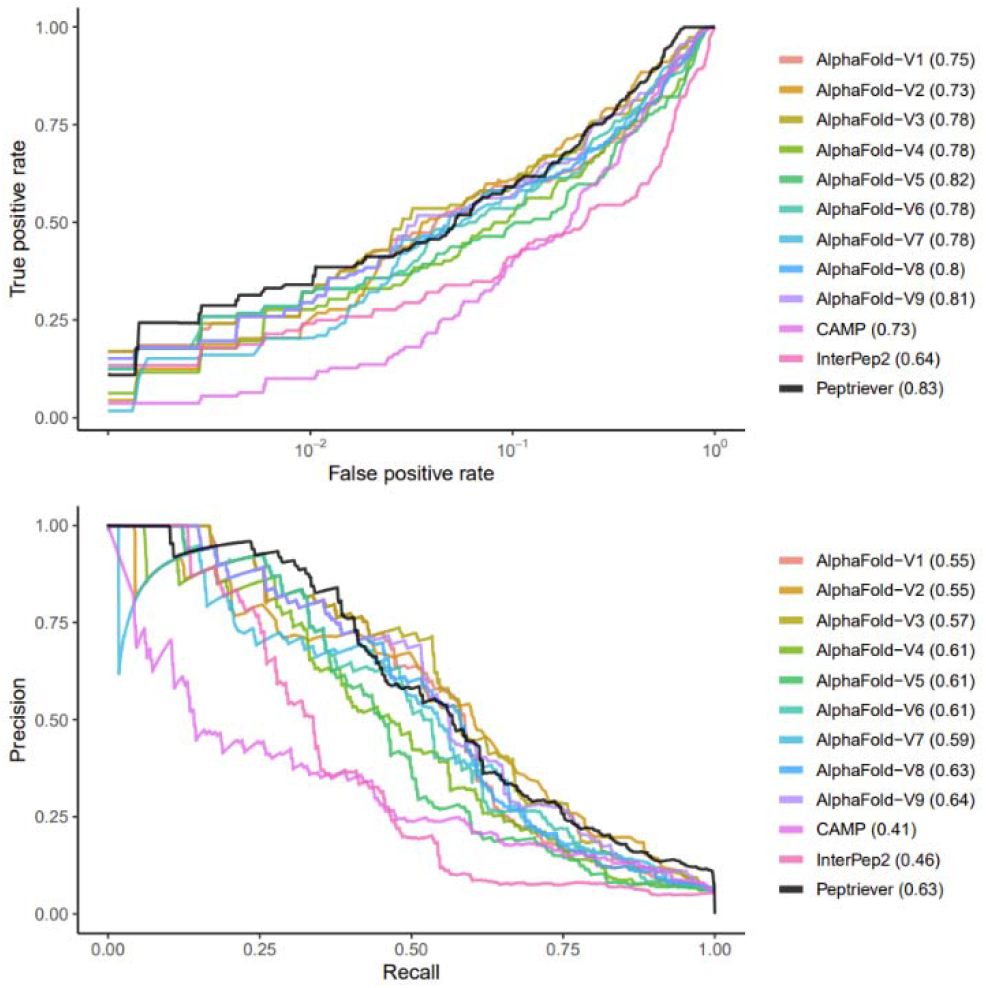
Performance measures for peptide-protein interaction prediction. (A) Receiver-operator curve (ROC) with the x-axis in log-scale. (B) Precision-recall (PR) curve. The values in the legends correspond to area under the ROC or PR curves. Curve coordinates from CAMP, InterPep2 and the various versions AlphaFold-Multimer were extracted from Johansson-Åkhe et al. using WebPlotDigitizer.

### 3.1. Application

The embeddings are stored in a vector database. Upon user input of a sequence, the database is searched for proteins with the highest binding probabilities, either across all species or a particular one as specified by the user.

### 3.2. Limitation

When training BERT-like models, sequences are adjusted to a fixed length through truncation or padding. We set the peptide and protein model input to 30 and 300 tokens, respectively. These lengths were chosen to encompass more than 90% of the proteins in the training set, ensuring that all the peptides were included. However, the performance may be suboptimal for proteins with longer sequences if important information is located towards the end, as it would be truncated.

## 4 Conclusion

Peptriever provides a user-friendly web interface, allowing users to input peptide sequences and retrieve protein binding scores across any number of proteins. The current version of Peptriever is limited to peptide search; however, in version II, we plan to add the feature to get binding peptides for the input protein sequences. Peptriever will be a useful tool for translational researchers to open new applications for peptide therapeutics and vaccine design.

